# Optimization of ^13^C stable isotope labeling for the study of tricarboxylic cycle intermediates in mouse models

**DOI:** 10.1101/2025.04.22.649802

**Authors:** Jarrod Laro, Monica Ness, Joseane Godinho, Randy Coats, Laura-Isobel McCall

## Abstract

The tricarboxylic acid cycle (TCA), also known as the Krebs Cycle or the citric acid cycle, is an essential metabolic pathway involved in energy production that is often impacted by disease, making it of key interest to identify effective, affordable, and simple ways to monitor the impact of disease on TCA metabolism. ^13^C-based stable isotope labeling is a useful technique to track pathway alterations in living hosts. However, infusion-based methodologies are slow and expensive despite achieving steady-state labeling. Bolus-based methods are cheaper, faster, and compatible with biohazardous models, but require optimization to achieve maximum labeling. Herein, we performed bolus-based stable isotope labeling experiments in mouse models to identify the optimal dosage amount, label administration length, fast length prior to label administration, ^13^C-labeled precursor, and route of administration for the TCA cycle in the esophagus, heart, kidney, liver, plasma, and proximal colon. ^13^C-glucose at a concentration of 4 mg/g administered via intraperitoneal injection followed by a 90 minute label incorporation period achieved the best overall TCA labeling. For most organs, a 3 hour fast prior to label administration improved labeling, but labeling in the heart was better with no fasting period, showcasing the need to optimize methodology on an organ-by-organ basis. We also identified that bolus administration of glucose provided little impact on metabolism compared to vehicle control. The experiments outlined here provide critical information for designing *in vivo* stable isotope labeling experiments for the study of the TCA cycle.

**Highlights:**

- A 90 minute waiting period following label administration provides best labeling
- Larger dosing provides better labeling with little impact on metabolism
- Fasting prior to label administration led to worse labeling in heart tissue
- Intraperitoneal dosing provides better incorporation than oral dosing
- ^13^C-glucose provides better label incorporation than ^13^C-lactate and ^13^C-pyruvate

## 1. Introduction

All elements have natural isotopes consisting of a fixed number of protons and variable number of neutrons which characterize the mass of the atom. In the instance of carbon, the three natural isotopes are ^12^C, ^13^C, and ^14^C. Approximately 99% of all carbons are ^12^C, 1% are ^13^C, and a negligible amount are ^14^C, meaning that almost all naturally occurring carbons are ^12^C. Techniques have been created that allow for the enrichment of molecules consisting of low abundance isotopes like ^13^C, making it possible to synthesize and isolate molecules consisting exclusively of ^13^C carbons. Despite the differences in masses between isotopes, ^12^C and ^13^C carbon molecules are treated identically by metabolic pathways, providing an interesting use case for ^13^C labeled molecules: ^13^C-labeled molecules can be administered to an organism and then metabolism can be measured in what is known as a stable isotope labeling experiment. In stable isotope labeling experiments, molecules which show a higher than normal fraction of ^13^C carbons following administration of a ^13^C precursor result from its metabolism, providing information on how the precursor is metabolised in the model of interest and at what rate. To provide a more concrete example, a naturally occurring 6-carbon molecule would be expected to have a little under 7% of molecules containing at least one ^13^C (i.e. a labeled fraction of 0.07) so molecules with significantly larger percentages of labeled molecules can be considered to be downstream products of the ^13^C precursor.

To observe how ^13^C-labeled precursors are incorporated into metabolism, they must first be administered to the test subject. The two most common methods for in vivo models are through either infusion or bolus injection[1]. Infusion methods involve infusing a constant flow of labeled precursors directly into the bloodstream, allowing for steady-state isotope labeling and a broader degree in labeling of downstream products than a bolus method[1]. To speed up the time required to achieve a steady state, sometimes a bolus dose is provided at the start of infusion[2–4]. Bolus methods involve dosing the subject with a large amount of labeled precursor and then waiting a fixed period time to allow incorporation into metabolism. While bolus injections can’t provide steady-state isotope labeling in vivo, their ease of use, speed, and lower costs and instrumental requirements make them more readily applicable in studies which require large sample sizes (n > 10)[5,6]. They are also more suitable for models of blood-borne pathogen infection, where setting up and maintaining an infusion pump could represent a biosafety liability. As a result, it is desirable to develop reliable, bolus-based steady isotope labeling protocols for infected mouse model studies.

The tricarboxylic acid cycle (TCA), also known as the citric acid cycle or Krebs cycle, is a critical biological pathway involved in electron transfer and generating precursors for other key biochemical pathways[7,8]. Due to its role in electron transfer, which is critical for energy production, it is particularly important in cell types that consume large amounts of ATP, such as skeletal muscle[9] and cardiac muscle[10]. Since it is a central part of cellular metabolism, many diseases involve some form of dysregulation of the TCA cycle, including Alzheimer’s disease[11], Huntington’s disease[12], Leigh syndrome[13],and some forms of cancer[14]. In the context of infectious disease, parasites like *Trypanaoma brucei* and *Plasmodium falciparum* are known to require TCA intermediates during periods of the growth cycles [15,16]. The prevalent role of the TCA cycle in several different diseases makes it a critical target for study, thus necessitating reliable, effective, and efficient methodology for studying its metabolic flux.

Herein, we discuss the approach we took to optimize ^13^C label administration methods suitable for the study of TCA intermediates (fumarate, succinate, alpha-ketoglutarate, malate, aconitate, and citrate/isocitrate) and some of their immediate precursors (glutamine, glutamate, G6P/F6P) by performing experiments covering common concerns around bolus label administration: dosage, the necessity and length of fast prior to label administration, the labeled precursor that provides best labeling, and the optimal route of label administration. Each study was performed across multiple sampling sites, to evaluate if different organs had different optimal parameters. Additionally, we investigated whether bolus glucose administration caused significant changes in the metabolism of our monitored metabolites when compared to mice provided with vehicle only.

## 2. Results/Discussion

### 2.1 Data Processing Optimization

Metabolism within live animals is complex and constantly occurring. While the mice are provided with a large bolus of fully labeled ^13^C_6_-glucose and fasted, ^12^C_6_-glucose is still naturally present in circulating blood, in stored glycogen, and in triglycerides. As a result, labeled metabolites are expected to range anywhere from partially labeled to fully labeled, meaning that it is not expected that every metabolite will be fully labeled. Additionally, the solvents used to extract TCA intermediates are fairly volatile, so samples could evaporate within the autosampler prior to liquid chromatography separation, causing concentrations to vary between samples due to differing liquid levels. To address these complications, we normalize the signal of each metabolite and its isotopologues by dividing the total intensity of the isotope by the sum of the intensity of all isotopologues of a metabolite (sometimes referred to as the Mass Distribution Vector, or MDV[17]). We then create two fractions for each metabolite: the unlabeled fraction consisting of only the M+0 isotope and the labeled fraction as the summation of all isotopes containing at least one labeled carbon.

To ensure the robustness of our data analytics pipeline and instrumental detection method, we mixed together yeast extract which was cultured with ^12^C-enriched media (^12^C yeast) and yeast extract which was cultured with ^13^C-enriched media (^13^C yeast) at 0%, 10%, 25%, 50%, 75%, 90%, and 100% fractions of ^13^C yeast to ^12^C yeast, and then tested to see if we could accurately identify the percentage of unlabeled vs labeled TCA intermediate fractions present in each sample. Ultimately, each measured metabolite fraction aligned closely with the expected fraction, demonstrating that this methodology can accurately differentiate between relative levels of ^13^C labeling (Figure 1).

**Figure 1.**
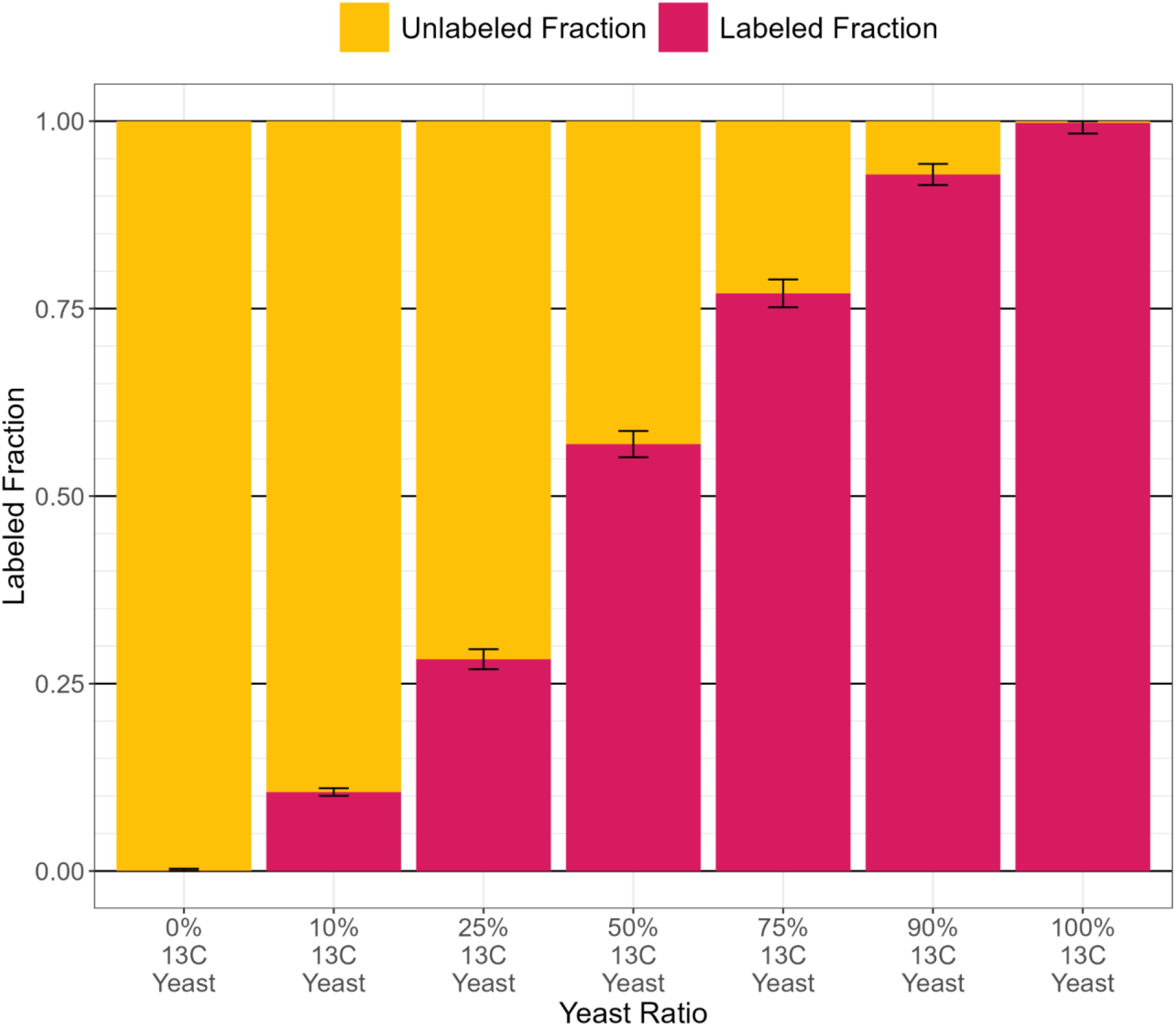
Relative fraction of unlabeled and labeled isotopes present in mixtures of ^12^C-enriched yeast and ^13^C-enriched yeast. Error bars represent the standard error of the mean. The labeled fraction represents the pooled fraction of glutamine, glutamate, fumarate, succinate, alpha-ketoglutarate, malate, fructose-6-phosphate, glucose-6-phosphate, aconitate, and citrate with at least one ^13^C. Error bars represent the standard error of the label fraction across the 10 measured metabolites for each sample.

### 2.2 Effects of Label Administration and Dosage

To achieve robust incorporation of ^13^C in TCA intermediates, it was important to identify an optimal dosage of ^13^C_6_-glucose as well as whether collecting each sample after a short waiting period following ^13^C_6_-glucose administration (15 minutes) or a longer period (90 minutes) would yield higher labeled fractions. In our first test cohort, we used a dosage of either 1 mg/g or 4 mg/g, which is near the maximum amount of glucose that can dissolve in 200 μL isotonic saline at room temperature for a 35 gram mouse. 15 minutes post-administration, plasma from the submandibular vein was collected (facial plasma). 90 minutes post-administration, plasma from veins in the abdomen (abdominal plasma) and the heart were collected. Aconitate, alpha-ketoglutarate, fumarate, glutamate, glutamine, and malate all showed significantly higher label fractions between the mice that received ^12^C_6_-glucose and mice that received ^13^C_6_-glucose, and significantly higher signal intensity between extraction blanks and ^13^C_6_-glucose samples (Figure S1). For both the 15 minute and 90 minute plasma samples, the mice that received 4 mg/g of ^13^C_6_-glucose showed a higher median labeled fraction of metabolites (Figure 2A; supplemental file 1). Between the 15 minute and 90 minute label administration for the 4 mg/g dosage, the 90 minute samples had a higher median labeled fraction. Focusing on the 4 mg/g 90 minute label administration samples, we examined if the labeled fraction from samples collected from mice that received ^13^C_6_-glucose differed from those which received ^12^C_6_-glucose. The only ^13^C_6_-glucose samples that did not have a higher median labeled fraction were the 1 mg/g heart samples (Figure 2B; supplemental file 1).

**Figure 2.**
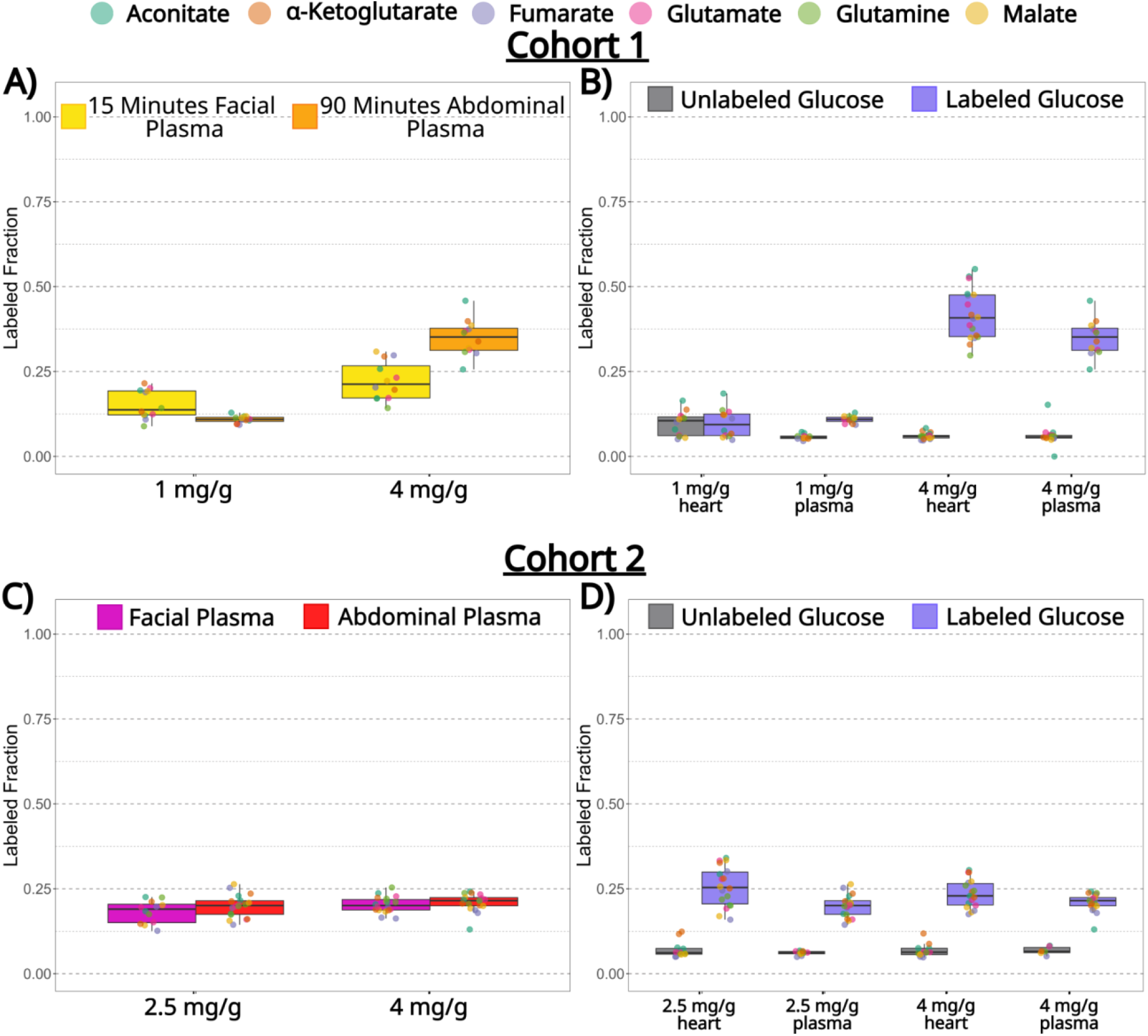
Effects of timing and dosage on labeling efficacy in mice. Individual measured metabolites are colored per the legend. A) Effects of collecting plasma 15 minutes after glucose administration vs 90 minutes after glucose administration. (1 mg/g n = 2 mice; 4 mg/g n = 3 mice). B) Comparison of labeled fractions 90 minutes after the administration of ^12^C_6_-glucose or ^13^C_6_-glucose in test cohort 1. (1 mg/g n = 2 mice; 4 mg/g n = 3 mice). C) Comparison of labeled fractions in facial plasma or abdominal plasma 90 minutes after administration of 2.5 mg/g or 4 mg/g of ^13^C_6_-glucose. (2.5 mg/g n = 3 mice; 4 mg/g n = 3 mice). D) Comparison of labeled fractions 90 minutes after the administration of ^12^C_6_-glucose or ^13^C_6_-glucose in test cohort 2. (2.5 mg/g n = 3 mice; 4 mg/g n = 3 mice).

Seeking to confirm that the non-terminally collected facial plasma used for the 15 minute time point is comparable to the terminally collected abdominal plasma and to identify if a middle-ground dose of 2.5 mg/g ^13^C_6_-glucose provided comparable labeling to 4 mg/g ^13^C_6_-glucose, we tested dosage effects on a second cohort of mice. In this cohort, facial plasma was collected 5 minutes prior to euthanasia and abdominal plasma collection to make them as comparable as possible. At both the 2.5 mg/g and 4 mg/g ^13^C_6_-glucose dosage, we saw no significant differences between the labeled fractions of the metabolites between the facial plasma and the abdominal plasma, highlighting a non-terminal way to monitor systemic ^13^C metabolism in mice (Figure 2C; supplemental file 1). In this cohort, both the 2.5 mg/g and 4 mg/g dosage of ^13^C_6_-glucose showed much higher labeled fraction than their ^12^C_6_-glucose counterpoints (Figure 2D; supplemental file 1). For future experiments, we decided to use 4 mg/g dosage after 90 minutes of administration since the 4 mg/g dosage in cohort 1 showed the highest labeled fraction, although 2.5 mg/g ^13^C_6_-glucose appeared to also provide acceptable labeling in both the heart and plasma.

### 2.3 Effects of Fasting Duration

Prior to the start of a labeling period, mice are typically fasted to allow for a neutral background in which labeling can be observed. This period is variable within the literature and can range from 3 hours to an overnight 12 hour fast all the way to a full 24 hour fast [18–21]. To assess the necessity of this fasting period, which inevitably causes a shift in metabolism, we either did not fast mice prior to glucose administration or fasted the mice for 3 hours prior to administration. All mice were still fasted during the 90 minute waiting period following glucose administration to prevent the introduction of new ^12^C_6_-glucose from their food source.

Generally, we observed that about half of observed metabolites in all organs showed no significant difference between non-fasted mice and mice fasted for 3 hours (Figure 3A). In kidney tissue, liver tissue, proximal colon tissue, and plasma, mice that were fasted for 3 hours prior to ^13^C bolus administration had higher labeled fractions in metabolites that were significantly different between the two groups (Figure 3A-D,F-G; Figure S2A). In spite of this, the heart showed no significant differences between the fasting durations in six of the measured molecules while the other 4 out of 10 had significantly higher labeled fractions in the mice that were not fasted (Figure 3A,E; Figure S2B). The esophagus showed no significant differences between mice that were fasted and mice that were not fasted (Figure 3A-B). While we saw differences in labeling levels based on the duration of the fast, it should be noted that all metabolites in all organs demonstrated significantly higher ^13^C labeling in mice that received ^13^C_6_-glucose than in mice that received ^12^C_6_-glucose.

**Figure 3.**
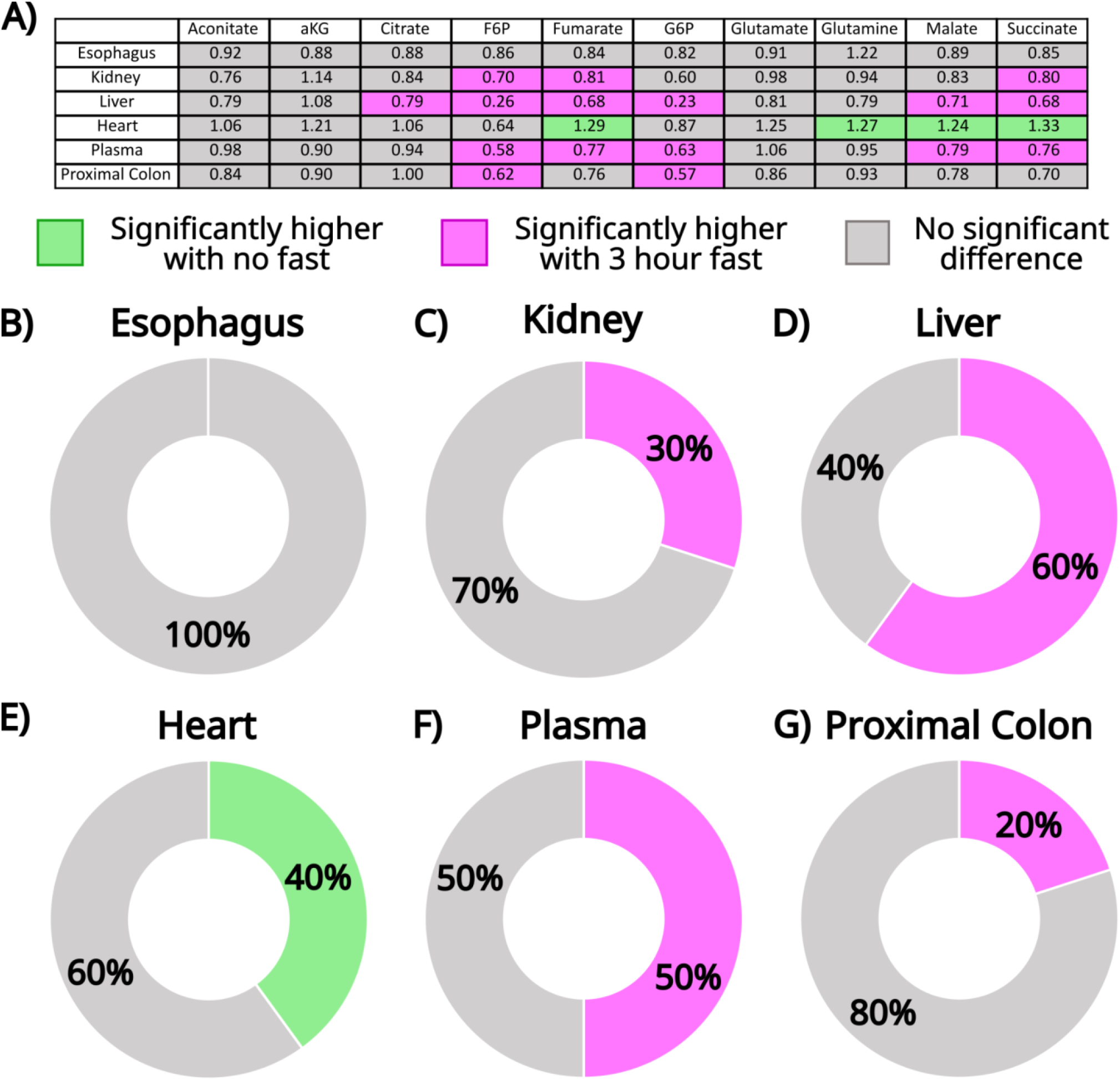
Fasting prior to label administration has differing impacts on label incorporation. A) Ratios of the median labeled fraction in mice that were not fasted prior to label administration to the median labeled fraction in mice that were fasted for three hours prior to label administration. Green fills are tissue and metabolite combinations that have significantly higher labeled fractions in the mice that were not fasted, while pink fills are tissue and metabolite combinations that have significantly higher labeled fractions in the mice that were fasted for 3 hours. (No fast, n = 5 mice; 3 hour fast, n = 5 mice). B-G) Percentage of the 10 measured metabolites showing a significant difference in labeled fraction between mice that were not fasted and mice that were fasted for 3 hours in B) esophagus, C) kidney, D) liver, E) heart, F) plasma, and G) proximal colon. (No fast, n = 5 mice; 3 hour fast, n = 5 mice).

Ultimately, we see that the benefits of fasting prior to label administration is somewhat tissue-dependent, possibly due to differences in ATP needs and TCA cycle turnover between organs[22]. The study these experiments were designed around focused primarily on energy-demanding tissues such as the heart and skeletal muscle, so we chose to delay fasting until after label administration. This approach was intended to maximize substrate availability and enhance labeled fraction uptake in these tissues.

### 2.4 Effects of Labeled Precursor and Administration Route

Having identified that a 90 minute label administration without a fasting period led to superior labeling of the heart and sufficient labeling in other organs, we next sought to identify if a different precursor or route of administration could provide even better labeling with these parameters. We tested fully labeled glucose (^13^C_6_), fully labeled lactate (^13^C_3_), and fully labeled pyruvate (^13^C_3_) administered via either an intraperitoneal bolus (IP) or an oral bolus. Both pyruvate and lactate are immediate precursors to the TCA cycle, so it was expected that they should serve as good alternatives to ^13^C_6_-glucose. All three precursors across both routes of administration showed significantly higher labeled fractions in mice that were provided with a fully labeled precursor compared to mice that received ^12^C_6_-glucose.

Overall, we observed significantly higher labeled fractions of each metabolite in the ^13^C_6_-glucose IP and ^13^C_6_-glucose oral route samples, when compared to their lactate and pyruvate counterparts (Figure 4; supplemental file 1). Between IP and orally administered ^13^C_6_-glucose, 40% of all measured metabolites had significantly larger labeled fractions in mice that received the labeled precursor by IP when compared to their oral counterparts (supplemental file 1). On average, the proximal colon had the largest difference between labeling routes, with a 1.52 fold increase in labeling via the IP route (Figure 5; supplemental file 1). The heart had the smallest average increase, with a fold change of 1.13 and only glucose-6-phosphate achieving statistical significance. Out of 48 measured metabolites (8 metabolites across 6 organs), only 6% of metabolites in ^13^C_3_-lactate-receiving mice (glucose-6-phosphate in the proximal colon; malate and succinate in the liver) and 8% of metabolites in ^13^C_3_-pyruvate-receiving mice (glucose-6-phosphate in the esophagus; glucose-6-phosphate and glutamate in plasma; malate in the liver) had significant differences in labeled fraction magnitude between IP and oral delivery (supplemental file 1). It should be noted that the dosage of both lactate (1.44 mg/g) and pyruvate (1.44 mg/g), although based on published literature values, was lower than glucose (4 mg/g) in this experiment[1,23]. However, even when comparing to mice that received the lower 2.5 mg/g ^13^C_6_-glucose dose, the average labeled fraction was still over 3.28 times lower in the heart for the mice that received ^13^C_3_-lactate or ^13^C_3_-pyruvate (compare to Figure 2D).

**Figure 4.**
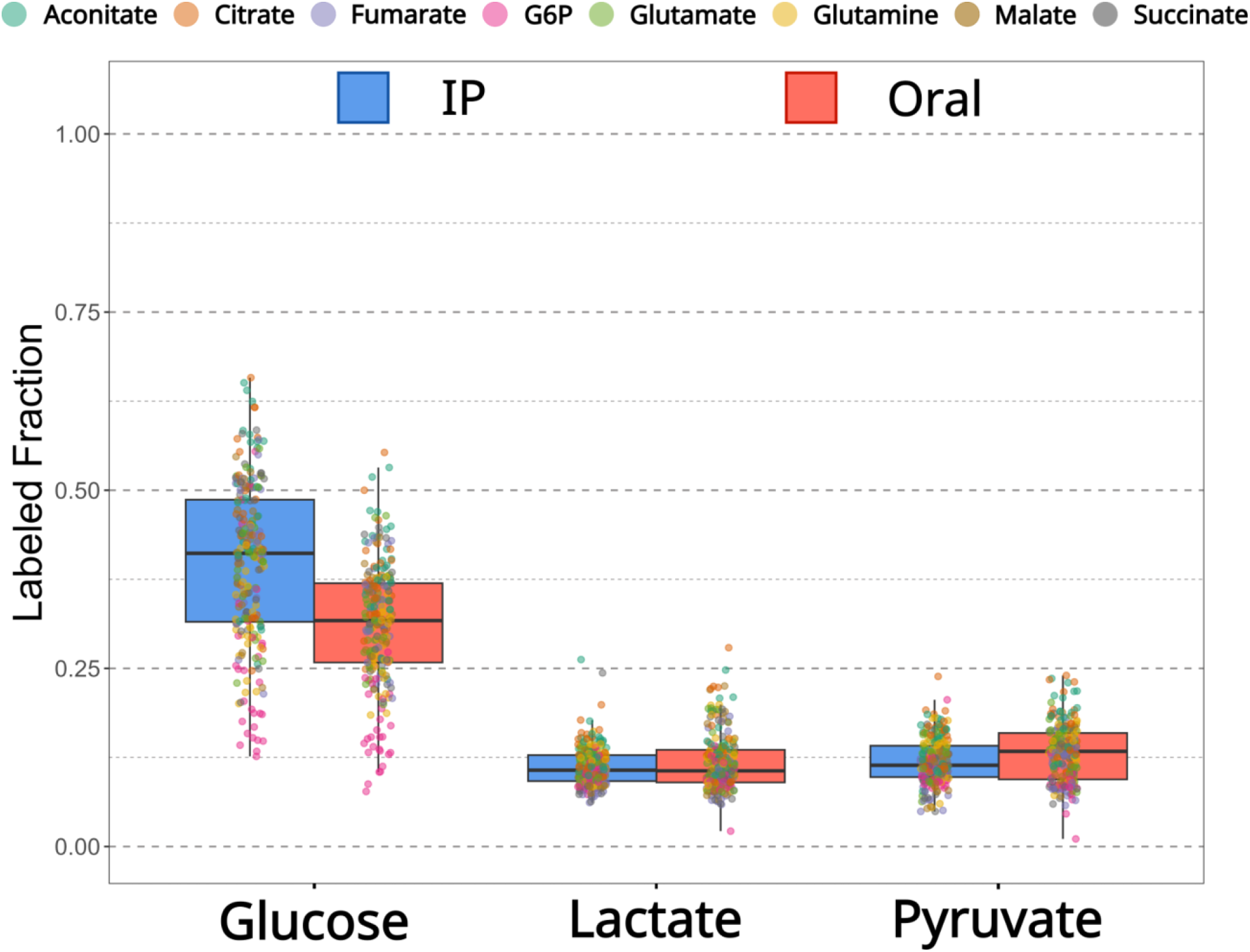
4 mg/g of ^13^C_6_-glucose delivered via IP injection provides the largest labeled fraction of target metabolites when compared to lactate and pyruvate via IP and oral administration routes. (all routes, all precursors n = 5 mice)

**Figure 5.**
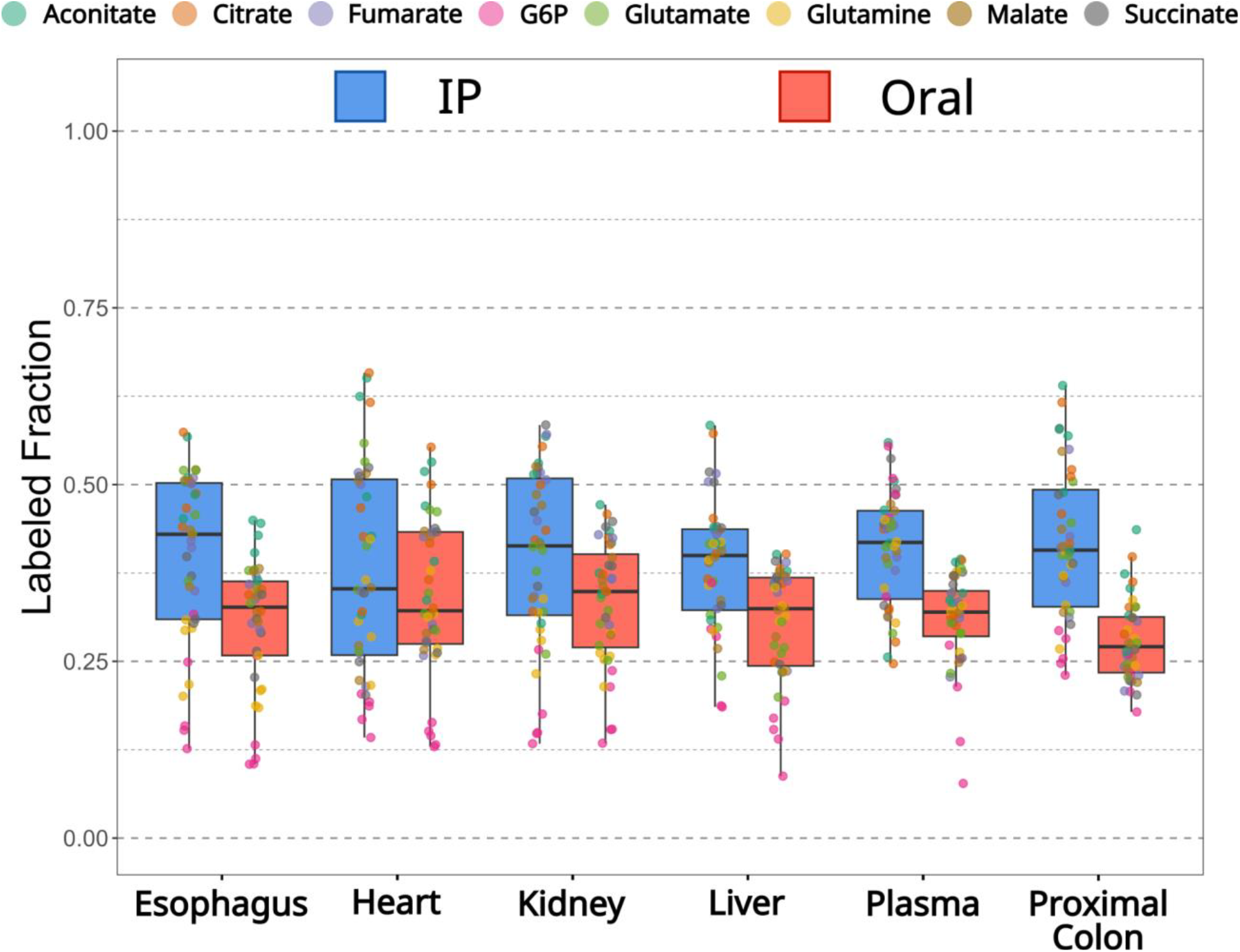
IP administration of ^13^C_6_-glucose provides larger labeled fractions than oral administration, particularly in the proximal colon. (both routes, n = 5 mice)

### 2.5 Effect of Glucose Bolus

A major concern with bolus-based labeling methods is that a sudden, large influx in precursor will cause unnatural shifts in metabolism [24]. We tested if the glucose bolus in both the oral and IP routes led to changes in raw signal intensity in downstream products of glucose metabolism when compared to vehicle without any glucose present. We observed little differences between the two groups via both the IP and oral routes (Figure 6). 4% had significant differences between the IP C12 mice and the IP vehicle mice, with only fumarate in the kidney showing a significantly higher signal intensity in the C12 mice compared to the vehicle while glutamate in the heart had significantly lower signal intensity (supplemental file 1). 10% had significant differences via the oral route (glucose-6-phosphate in the kidney; glutamate in the liver; glutamine in the liver, proximal colon, and kidney), but all had significantly lower signal intensities compared to the vehicle (supplemental file 1). It would be expected that a bolus of glucose would cause an increase in TCA activity to metabolize the glucose bolus and re-achieve metabolic equilibrium, so the decrease in signal intensity indicates that these observed differences are not an artefact induced by the bolus precursor amounts.

**Figure 6.**
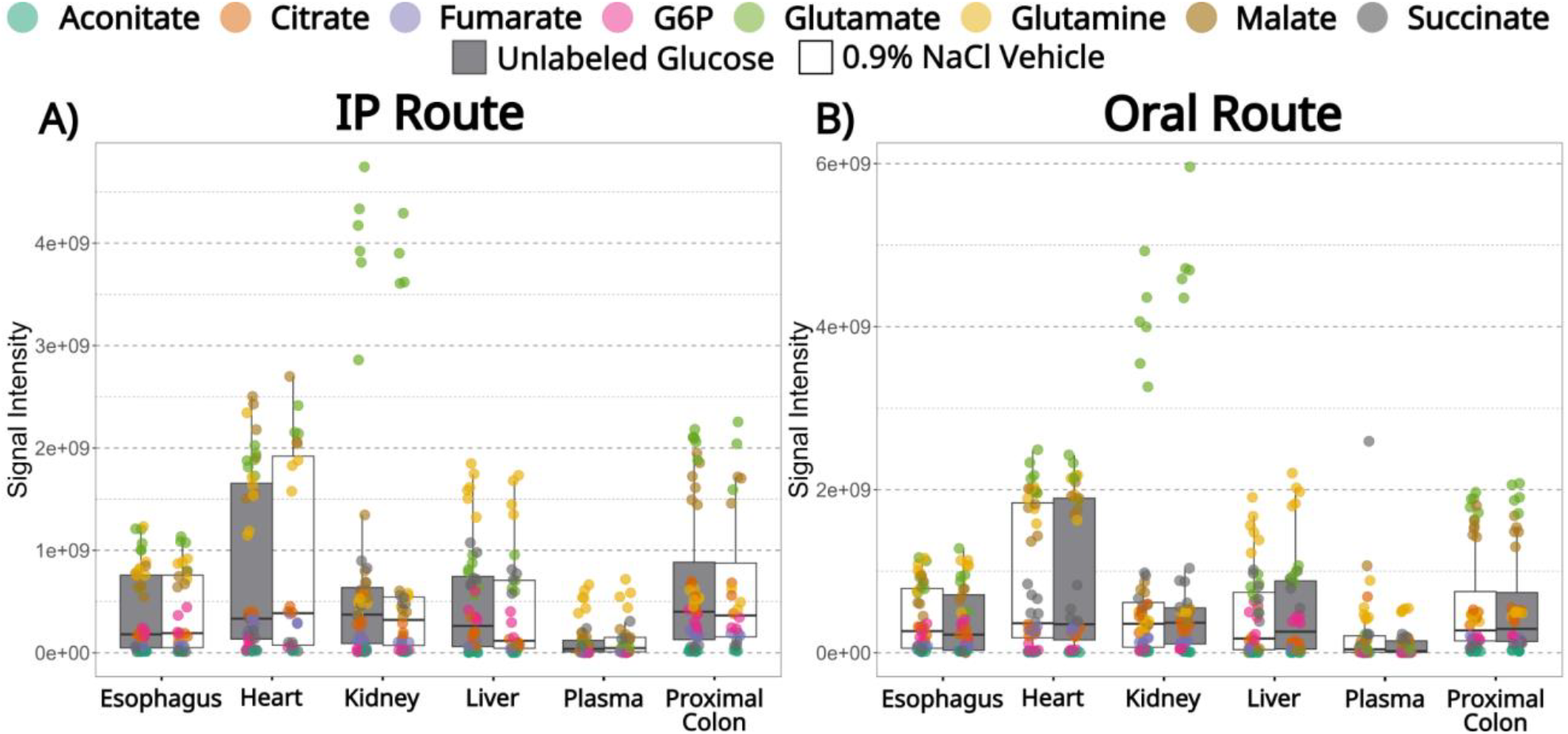
^12^C_6_-glucose administration shows little effects on TCA metabolism via A) IP administration and B) oral administration

## 3. Conclusion

Here, we show that bolus administration of ^13^C_6_-glucose provides sufficient labeling of TCA intermediates and immediate precursors. We found that 4 mg/g dosage provided the largest labeled fractions, although as little as 1 mg/g was sufficient for labeling in the plasma and 2.5 mg/g was sufficient for labeling in the heart. Fasting prior to label administration, a standard practice in steady isotope labeling, did improve labeling efficiency in most organs. However, it led to worse labeling in the heart tissue after a 90 minute label administration period, demonstrating that it might not necessarily be best practice for all steady isotope labeling studies and highlighting the importance of validating a method for the organ of highest interest. Administration of glucose via IP injection provided better labeling than oral administration, and the use of ^13^C_6_-glucose outperformed fully labeled lactate and pyruvate, although at higher doses relative to the lactate and pyruvate. Finally, we saw that bolus administration of glucose caused little shift in metabolism in the context of the TCA and some of its immediate precursors. While the optimization that occurred in this study focused specifically on the TCA cycle and might not apply directly to the study of other metabolic pathways, the experiments performed here highlight the necessity for tracer administration optimization prior to beginning a steady isotope labeling experiment, to ensure the methodology employed provides sufficient labeling for the organ and pathway of choice. This method should be of use for investigators who are unable to use infusion for logistical reasons, such as large treatment cohorts or infectious systems.

## 4. Methods

### 4.1 Data Processing Optimization and Labeled Yeast Samples

^13^C yeast [Metabolite Yeast Extract (U-^13^C, 99%); ISO1] and ^12^C yeast [Metabolite Yeast Extract (unlabeled); ISO1-UNL] were purchased from Cambridge Isotope Laboratories. Yeast extracts were combined at 0%, 10%, 25%, 50%, 75%, 90%, and 100% fractions of ^13^C yeast to ^12^C yeast, sonicated for 5 minutes, and then centrifuged for 10 mins at 12,000 RPM. Each combination of yeast extracts was then placed in its own well on Level 2 96-well round bottom plates (Thermo Scientific).

### 4.2 Mouse models

#### 4.2.1 Animal Ethics Statement

All experiments were performed under University of Oklahoma Institutional Animal Care and Use Committee approved protocol number R20-027.

#### 4.2.2 Effects of Label Administration and Dosage

##### 4.2.2.1 Cohort 1

5-week-old male Swiss Webster mice were purchased from Jackson labs. Mice were IP injected with 1 mg/g of ^13^C_6_-glucose (n = 2), 4 mg/g of ^13^C_6_-glucose (n = 3), 1 mg/g of ^12^C_6_-glucose (n = 2), or 4 mg/g of ^12^C_6_-glucose (n = 3). Dosage mass was determined the day prior to the experiment by averaging the mass of the mice in each experimental group. All solutions were administered in 200 μL of 0.9% NaCl isotonic saline solution. Access to food was removed upon label administration. After 15 minutes, a facial bleed was performed on each mouse. After 90 minutes, mice were euthanized with isoflurane and both blood and heart tissue were collected for LC-MS analysis. Plasma was collected from all blood samples by centrifugation at 12,000 RPM for 10 minutes. All samples were snap frozen in liquid nitrogen upon collection.

##### 4.2.2.2 Cohort 2

5-week-old male Swiss Webster mice were purchased from Jackson labs.^13^C_6_-glucose was purchased from Cambridge Isotope Laboratories. Mice were IP injected with 2.5 mg/g of ^13^C_6_-glucose (n = 3), 4 mg/g of ^13^C_6_-glucose (n = 3), 2.5 mg/g of ^12^C_6_-glucose (n = 2), or 4 mg/g of ^12^C_6_-glucose (n = 2). Dosage mass was determined the day prior to the experiment by averaging the mass of the mice in each experimental group. All solutions were administered in 200 μL 0.9% NaCl isotonic saline solution. 85 minutes post-label administration, a facial bleed was performed on each mouse. 90 minutes post-label administration, mice were euthanized with isoflurane and both plasma and heart tissue were collected for LC-MS analysis. Plasma was collected from all blood samples by centrifugation at 12,000 RPM for 10 minutes. All samples were snap frozen in liquid nitrogen upon collection.

#### 4.2.3 Effects of Fasting Duration

5-week-old male Swiss Webster mice were purchased from The Jackson Laboratory. Mice were either fasted for 3 hours prior to label administration and continued to be fasted until euthanasia (4.5 hours total; n = 8) or were fasted upon label administration (1.5 hours total; n = 7). Each mouse received either 4 mg/g ^12^C_6_-glucose (n = 2 for 1.5 hr fast; n = 3 for 4.5 hr fast) or 4 mg/g ^13^C_6_-glucose by I.P. injection (n = 5 for both fasting durations). 90 minutes post-label administration, mice were euthanized with isoflurane and plasma, heart tissue, esophageal tissue, proximal colon tissue, kidney tissue, and liver tissue were collected for LC-MS analysis. All samples were snap frozen in liquid nitrogen upon collection.

#### 4.2.4 Effects of Labeled Precursor and Administration Route

5-week-old male Swiss Webster mice were purchased from Jackson labs. ^13^C_6_-glucose, ^13^C_3_-sodium lactate, and ^13^C_3_-sodium pyruvate were purchased from Cambridge Isotope Laboratories. Mice were not fasted until precursor administration. Mice were split into one of two cohorts: either a cohort receiving precursor via IP injection or a cohort receiving precursor via oral gavage. In the IP cohort, mice either received 4 mg/g of glucose (^12^C_6_ n = 2; ^13^C_6_ n = 5), 1.44 mg/g of lactate (^12^C_3_ n = 2; ^13^C_3_ n = 5), 1.44 mg/g of pyruvate (^12^C_3_ n = 2; ^13^C_3_ n = 5), or vehicle (n = 4). Similarly in the oral gavage cohort, mice either received 4 mg/g of glucose (^12^C_6_ n = 2; ^13^C_6_ n = 5), 1.44 mg/g of lactate (^12^C_3_ n = 2; ^13^C_3_ n = 5), 1.44 mg/g of pyruvate (^12^C_3_ n = 2; ^13^C_3_ n = 5), or vehicle (n = 5). All solutions were administered in 200 μL 0.9% NaCl isotonic saline solution. Lactate and pyruvate dosage amounts were based on previously published studies[1,23].

90 minutes post-label administration, mice were euthanized with isoflurane, and plasma, heart tissue, esophageal tissue, proximal colon tissue, kidney tissue, and liver tissue were collected for LC-MS analysis. All samples were snap frozen in liquid nitrogen upon collection.

### 4.3 Metabolite Extraction and LC-MS Analysis

To account for differences in tissue weights and collected plasma volumes, extraction solvent volumes were adjusted relative to the weight of the sample. Samples were extracted in ice cold LC-MS grade 80:20 ACN:Water + 2 μM sulfachloropyridazine to serve as an internal standard. A 3 mm tungsten carbide bead was added to each samples and then homogenized for 5 min at 5 Hz using a TissueLyser II (Qiagen). Afterwards, samples were sonicated for 5 minutes followed by centrifugation for 10 mins at 12,000 RPM. 120 μL of supernatant was plated into a Level 2 96-well round bottom plates (Thermo Scientific) and then dried down in a SpeedVac overnight.

Samples were resuspended in 150 μL LC-MS grade 80:20 ACN:Water + 2 μM sulfadimethoxine, sonicated for 5 minutes, and centrifuged for 10 minutes at 4,000 RPM. 120 μL were transferred to an Level 2 96-well round bottom plates. For each tissue type, 5 μL of each tissue sample were pooled into a single well to serve as a quality control during the course of the LC-MS analysis. Samples were analyzed using a Thermo Scientific Vanquish UHPLC equipped with a SeQuant ZIC-pHILIC 5 µm column (150 mm x 2.1 mm, Sigma). Mobile phase A was LC-MS grade water (Fisher Optima) containing 20 mM ammonium carbonate adjusted to pH 9.4 using ammonium hydroxide. Mobile phase B was unmodified LC-MS grade acetonitrile. An 18 minute gradient was used with the parameters listed in Table S1. Liquid chromatography gradient parameters and mobile phases were validated using pure TCA standards. A Thermo Scientific Q Exactive Plus high resolution orbitrap mass spectrometer was used in negative ionization mode to collect untargeted MS/MS data (exact parameters as in Table S2).

### 4.4 Data analysis

Raw chromatograms were converted into peak areas using Xcalibur 4.7. For metabolites of interest, the raw peak areas were compared between the labeled samples and process blank samples to confirm that signal was significantly higher in both unlabeled peak area and pooled labeled peak area. Afterwards, unlabeled and labeled peak areas were converted into fractions by dividing the peak area of each isotopologue by the total ion current of all possible isotopes. Metabolites of interest were then compared between samples administered with a ^12^C precursor and samples administered with a ^13^C precursor, to confirm that the ^13^C samples had significantly higher labeled fractions. Any molecule of interest that did not meet either of these criteria were excluded from further analysis. All statistical analysis was performed using R Studio 4.1.

## Author Contributions

Conceptualization, J.L, L-I.M.; Data Curation, J.L.; Formal Analysis, J.L.; Funding Acquisition, L-I.M; Investigation, J.L., M.N., J.G., and R.C.; Methodology, J.L.; Project Administration, L-I.M.; Software, J.L.; Resources, L-I.M.; Supervision, L-I.M.; Visualization, J.L.; Writing – Original Draft, J.L.; Writing – Review & Editing, J.L., M.N., J.G., R.C., and L-I.M.

## Acknowledgements

Research reported in this publication was supported by the National Institute Of Allergy And Infectious Diseases of the National Institutes of Health under Award Number R01AI170605. The content is solely the responsibility of the authors and does not necessarily represent the official views of the National Institutes of Health.

## Supplementary Data

**Supplemental Figure 1.**
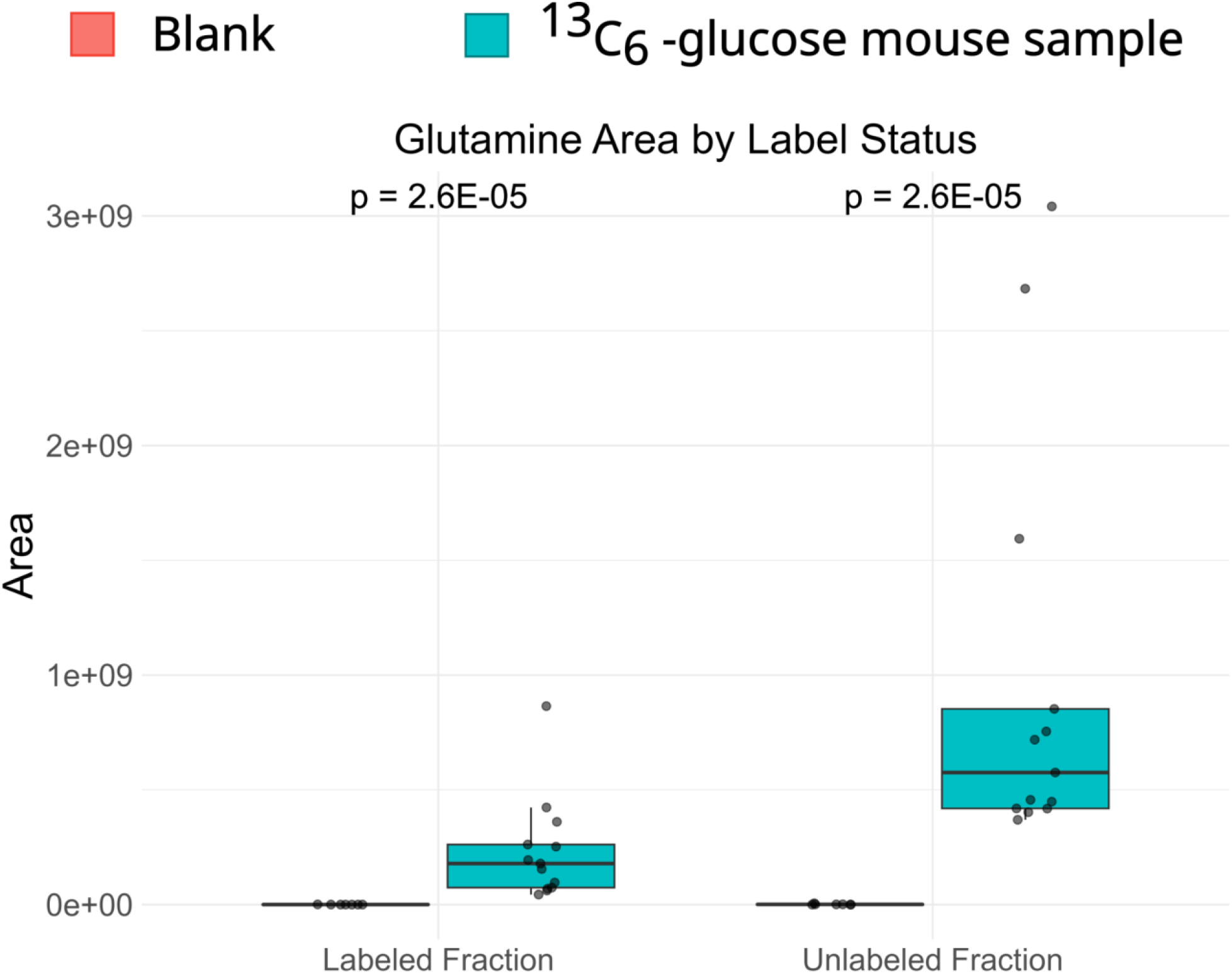
Representative example of the differences between ^13^C labeled samples and process blanks (Blanks, n = 7; ^13^C labeled samples, n = 13 samples).

**Supplemental Figure 2.**
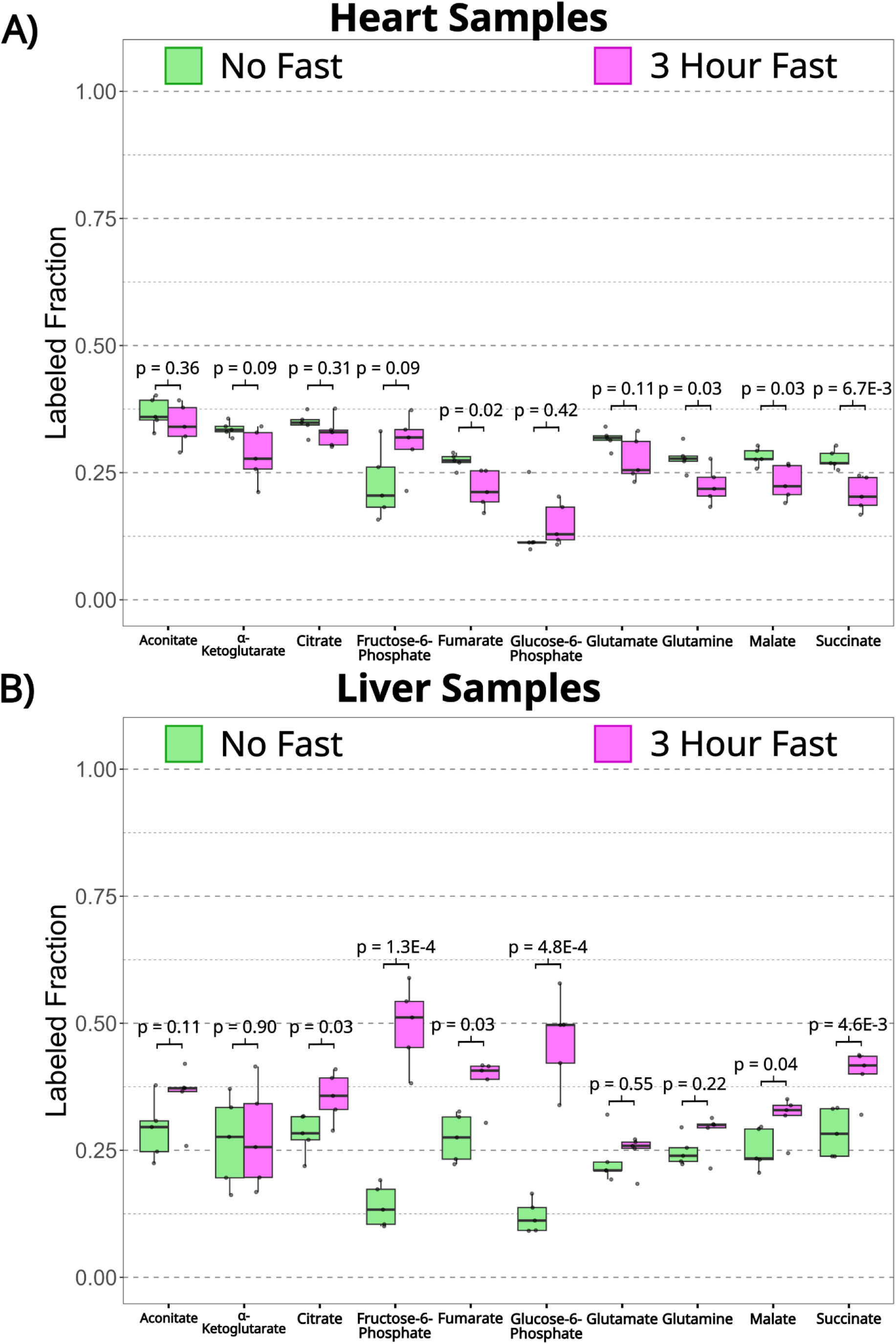
Labeled fractions of all measured metabolites for mice that were not fasted and mice that were fasted for 3 hours (n = 5 mice). A) Heart samples. B) Liver samples.

Supplemental File 1 - Full lists of p-values and fold changes for the effect of label administration, effects of precursor and administration route, and effects of glucose bolus.

**Supplemental Table 1.** Liquid chromatography parameters.

| **Time [min]** | **Flow**<br>**[mL/min]** | **%B** | **Curve** |
| --- | --- | --- | --- |
| 0.000 | 0.250 | 90.0 | 5 |
| 10.000 | 0.250 | 30.0 | 5 |
| 12.000 | 0.250 | 30.0 | 5 |
| 12.666 | 0.250 | 90.0 | 5 |
| 18.000 | 0.250 | 90.0 | 5 |

**Supplemental Table 2.** Mass spectrometer parameters.

|  |  |
| --- | --- |
| Run time | 0 to 12.7 min |
| Polarity | Negative |
| Default charge state | 1 |
| **Full MS** |  |
| Resolution | 70,000 |
| AGC target | 3e6 |
| Maximum IT | 246 ms |
| Scan range | 80 to 1200 *m/z* |
| **dd-MS2** |  |
| Resolution | 17,500 |
| AGC target | 1e5 |
| Maximum IT | 54 ms |
| Loop count | 5 |
| TopN | 5 |
| Isolation window | 1.0 *m/z* |
| (N)ce/stepped (N)CE | NCE: 20, 40, 60 |
| **dd Settings** |  |
| Minimum AGC | 8e3 |
| Intensity Threshold | 1.5e5 |
| Peptide match | Preferred |
| Exclude isotopes | Off |
| Dynamic exclusion | 10.0 s |

## Data availability

Data has been deposited into MassIVE, massive.ucsd.edu, accession number MSV000097457. All code used for the statistical analysis in this project can be found at https://github.com/jarrodRoachChem/C13Methods

## References

[1] B. Faubert, A. Tasdogan, S.J. Morrison, T.P. Mathews, R.J. DeBerardinis, Stable isotope tracing to assess tumor metabolism in vivo, Nat. Protoc. 16 (2021) 5123–5145. 10.1038/s41596-021-00605-2.

[2] M. Lai, B. Lanz, C. Poitry-Yamate, J.F. Romero, C.M. Berset, C. Cudalbu, R. Gruetter, In vivo 13C MRS in the mouse brain at 14.1 Tesla and metabolic flux quantification under infusion of [1,6-13C2]glucose, J. Cereb. Blood Flow Metab. 38 (2018) 1701–1714. 10.1177/0271678X17734101.

[3] S. Li, Y. Zhang, M. Ferraris Araneta, Y. Xiang, C. Johnson, R.B. Innis, J. Shen, In vivo detection of 13C isotopomer turnover in the human brain by sequential infusion of 13C labeled substrates, J. Magn. Reson. 218 (2012) 16–21. 10.1016/j.jmr.2012.03.012.

[4] D.K. Deelchand, C. Nelson, A.A. Shestov, K. Uğurbil, P.-G. Henry, Simultaneous measurement of neuronal and glial metabolism in rat brain in vivo using co-infusion of [1,6-13C2]glucose and [1,2-13C2]acetate, J. Magn. Reson. 196 (2009) 157–163. 10.1016/j.jmr.2008.11.001.

[5] A. Moreno, S. Blüml, J.H. Hwang, B.D. Ross, Alternative 1-(13)C glucose infusion protocols for clinical (13)C MRS examinations of the brain, Magn. Reson. Med. 46 (2001) 39–48. 10.1002/mrm.1158.

[6] S.-P. Wang, D. Zhou, Z. Yao, S. Satapati, Y. Chen, N.A. Daurio, A. Petrov, X. Shen, D. Metzger, W. Yin, A.R. Nawrocki, G.J. Eiermann, J. Hwa, C. Fancourt, C. Miller, K. Herath, T.P. Roddy, D. Slipetz, M.D. Erion, S.F. Previs, D.E. Kelley, Quantifying rates of glucose production in vivo following an intraperitoneal tracer bolus, Am. J. Physiol. Endocrinol. Metab. 311 (2016) E911–E921. 10.1152/ajpendo.00182.2016.

[7] P.K. Arnold, L.W.S. Finley, Regulation and function of the mammalian tricarboxylic acid cycle, J. Biol. Chem. 299 (2023) 102838. 10.1016/j.jbc.2022.102838.

[8] J.M. Borkum, The tricarboxylic acid cycle as a central regulator of the rate of aging: Implications for metabolic interventions, Adv. Biol. (Weinh.) 7 (2023) e2300095. 10.1002/adbi.202300095.

[9] M.J. Gibala, D.A. MacLean, T.E. Graham, B. Saltin, Tricarboxylic acid cycle intermediate pool size and estimated cycle flux in human muscle during exercise, Am. J. Physiol. Endocrinol. Metab. 275 (1998) E235–E242. 10.1152/ajpendo.1998.275.2.E235.

[10] S. Veera, F. Tang, Y. Mourad, S. Kim, T. Liu, H. Li, Y. Wang, J.S. Warren, J. Park, C. Van, J. Sadoshima, S.-I. Oka, A transcriptional regulatory mechanism of genes in the tricarboxylic acid cycle in the heart, J. Biol. Chem. 300 (2024) 107677. 10.1016/j.jbc.2024.107677.

[11] D. Jia, F. Wang, H. Yu, Systemic alterations of tricarboxylic acid cycle enzymes in Alzheimer’s disease, Front. Neurosci. 17 (2023) 1206688. 10.3389/fnins.2023.1206688.

[12] N.N. Naseri, H. Xu, J. Bonica, J.P.G. Vonsattel, E.P. Cortes, L.C. Park, J. Arjomand, G.E. Gibson, Abnormalities in the tricarboxylic Acid cycle in Huntington disease and in a Huntington disease mouse model, J. Neuropathol. Exp. Neurol. 74 (2015) 527–537. 10.1097/NEN.0000000000000197.

[13] J.-J. Brière, J. Favier, A.-P. Gimenez-Roqueplo, P. Rustin, Tricarboxylic acid cycle dysfunction as a cause of human diseases and tumor formation, Am. J. Physiol. Cell Physiol. 291 (2006) C1114–20. 10.1152/ajpcell.00216.2006.

[14] J. Eniafe, S. Jiang, The functional roles of TCA cycle metabolites in cancer, Oncogene 40 (2021) 3351–3363. 10.1038/s41388-020-01639-8.

[15] O. Villafraz, M. Biran, E. Pineda, N. Plazolles, E. Cahoreau, R. Ornitz Oliveira Souza, M. Thonnus, S. Allmann, E. Tetaud, L. Rivière, A.M. Silber, M.P. Barrett, A. Zíková, M. Boshart, J.-C. Portais, F. Bringaud, Procyclic trypanosomes recycle glucose catabolites and TCA cycle intermediates to stimulate growth in the presence of physiological amounts of proline, PLoS Pathog. 17 (2021) e1009204. 10.1371/journal.ppat.1009204.

[16] J.I. MacRae, M.W. Dixon, M.K. Dearnley, H.H. Chua, J.M. Chambers, S. Kenny, I. Bottova, L. Tilley, M.J. McConville, Mitochondrial metabolism of sexual and asexual blood stages of the malaria parasite Plasmodium falciparum, BMC Biol. 11 (2013) 67. 10.1186/1741-7007-11-67.

[17] J. Fernández-García, P. Altea-Manzano, E. Pranzini, S.-M. Fendt, Stable isotopes for tracing mammalian-cell metabolism in vivo, Trends Biochem. Sci. 45 (2020) 185–201. 10.1016/j.tibs.2019.12.002.

[18] M. Lopes, K. Brejchova, M. Riecan, M. Novakova, M. Rossmeisl, T. Cajka, O. Kuda, Metabolomics atlas of oral 13C-glucose tolerance test in mice, Cell Rep. 37 (2021) 109833. 10.1016/j.celrep.2021.109833.

[19] M. Mishkovsky, B. Anderson, M. Karlsson, M.H. Lerche, A.D. Sherry, R. Gruetter, Z. Kovacs, A. Comment, Measuring glucose cerebral metabolism in the healthy mouse using hyperpolarized 13C magnetic resonance, Sci. Rep. 7 (2017) 11719. 10.1038/s41598-017-12086-z.

[20] P. Lee, W. Leong, T. Tan, M. Lim, W. Han, G.K. Radda, In vivo hyperpolarized carbon-13 magnetic resonance spectroscopy reveals increased pyruvate carboxylase flux in an insulin-resistant mouse model, Hepatology 57 (2013) 515–524. 10.1002/hep.26028.

[21] M. Grima-Reyes, A. Martinez-Turtos, I. Abramovich, E. Gottlieb, J. Chiche, J.-E. Ricci, Physiological impact of in vivo stable isotope tracing on cancer metabolism, Mol. Metab. 53 (2021) 101294. 10.1016/j.molmet.2021.101294.

[22] B.Y. Nguyen, A. Ruiz-Velasco, T. Bui, L. Collins, X. Wang, W. Liu, Mitochondrial function in the heart: the insight into mechanisms and therapeutic potentials: Mitochondrial dysfunction and heart diseases, Br. J. Pharmacol. 176 (2019) 4302–4318. 10.1111/bph.14431.

[23] Z. Chen, B. Bordieanu, R. Kesavan, N.P. Lesner, S.S.K. Venigalla, S.D. Shelton, R.J. DeBerardinis, P. Mishra, Lactate metabolism is essential in early-onset mitochondrial myopathy, Sci. Adv. 9 (2023) eadd3216. 10.1126/sciadv.add3216.

[24] R.D. Sheldon, E.H. Ma, L.M. DeCamp, K.S. Williams, R.G. Jones, Interrogating in vivo T-cell metabolism in mice using stable isotope labeling metabolomics and rapid cell sorting, Nat. Protoc. 16 (2021) 4494–4521. 10.1038/s41596-021-00586-2.

